# DEX: an amino acid exchangeability measure for codon substitution modelling and selection inference

**DOI:** 10.64898/2026.03.09.710665

**Authors:** Gavin M. Douglas, Louis-Marie Bobay

## Abstract

Physicochemically similar amino acids undergo more frequent substitutions compared to dissimilar amino acid pairs. Despite their clear potential, amino acid similarity matrices remain underused for certain molecular evolution applications. One key potential application that is understudied is in quantifying the strength of natural selection based on amino acid substitution patterns. This is partially due to the high number of proposed amino acid distance measures and the lack of agreement on which are most accurate. In this study, we assessed the performance of 30 amino acid distance measures, including a new amino acid distance measure we developed based on recent deep mutational scanning data. We compared these measures across codon substitution models fit to alignments spanning *Streptococcus*, *Drosophila*, and mammalian lineages, as well as segregating variants across *Escherichia coli* strains and human genotypes. We further constructed consensus matrices from combinations of top-performing measures in this analysis using the DISTATIS approach and retested these matrices. Our results show that experimentally-derived measures, particularly our new measure, DMS-EX and the existing experimental exchangeability measure, best fit codon substitution patterns across diverse lineages. We found that a consensus measure based on these two approaches, which we named DEX, performed best overall. We also explored the value of asymmetric exchangeabilities in DMS-EX for predicting allele frequencies of replacement polymorphisms across diverse lineages, including when conditioned on buried vs. exposed sites. Overall, we provide a systematic comparison of the performance of existing measures. The amino acid distance measures we introduce constitute a substantial improvement for exploring novel methods for quantifying the strength of natural selection and for providing improved baselines for future benchmarking approaches.

**Significance:** Protein-coding genes have long been a focus for researchers studying the strength and direction of selection. By studying non-synonymous substitutions, those that change amino acids, it is possible to estimate the relative strength of selection. Despite widespread interest in such approaches, information on which amino acids are exchanged is underused in most molecular evolution applications. This is partly because many different measures exist for quantifying amino acid distances, particularly those based on physicochemical properties. A newer class of amino acid distance measures is derived from deep mutational scanning datasets, where virtually every possible substitution is tested for its impact on protein function. We characterised and compared 30 amino acid distance measures, including a novel measure based on deep mutational scanning data. We highlight differences in how well these measures fit real substitution data. Overall, we find that DEX, which is a consensus of our new measure and an existing experimental exchangeability measure, performed best in these models. This work will serve as the basis for future improved methods for inferring selection efficacy from protein-coding alignments.

## Introduction

Amino acid substitutions have long been known to be more frequent between physicochemically similar amino acids relative to dissimilar pairs (Zuckerkandl and Pauling 1965; Dayhoff et al. 1978; Graur 1985). These biases reflect the action of natural selection for the conservation of protein folding and function. Nonetheless, amino acid distances are often ignored or underused when testing for selection based on non-synonymous substitution patterns. One key reason why amino acid distances remain underused in this context is that many contrasting measures have been proposed to summarize physicochemical properties and other characteristics. A comparison of amino acid distance measures is needed to assess which are most generalizable and useful for molecular evolution methods.

A standard test for natural selection acting on protein-coding DNA alignments is based on the ratio of non-synonymous to synonymous substitution rates (d_N_/d_S_) (Muse 1996). Early forms of this test explicitly incorporated amino acid distances (Miyata and Yasunaga 1980; Li et al. 1985; Goldman and Yang 1994), but the most common workflows now treat all amino acid substitutions as equally likely. These tests are based on a codon substitution model where the independent action of varying mutation rate for specific nucleotide changes and selection acting on non-synonymous changes can be modelled separately. A similar approach is the ratio of radical to conservative amino acid substitutions (d_R_/d_C_) (Hughes et al. 1990; Zhang 2000). This framework is based on binning amino acid substitutions into those that occur between highly dissimilar amino acids (radical) and those that occur between similar amino acids (conservative). Although d_R_/d_C_ is a valuable approach, particularly when corrected for compositional biases (Dagan et al. 2002; Weber and Whelan 2019), it relies on the simplistic binary categorization of substitutions despite continuous distances having the potential of better capturing the relative impact of selection acting on the different types of substitutions (Braun 2018; James and Lascoux 2025). Amino acid exchangeability has also been considered in other extensions of d_N_/d_S_ models, which focus on the most exchangeable amino acids when computing d_N_, rather than all amino acids (Tang and Wu 2006). This approach showed increased power for detecting positive selection, but it is based on empirically observed substitution patterns rather than independent data, which risks circularity. All of these approaches, whether based on explicitly incorporating amino acid distances, or binning substitutions into groups, require accurate amino acid distance matrices. We hypothesize that methods incorporating continuous amino acid measures, which have been understudied in molecular evolution selection tests, could better estimate selection efficacy. However, understanding the relative utility of different amino acid distance measures is a necessary first step, and is our focus herein.

Two commonly used amino acid distance measures in molecular evolution are Grantham’s (Grantham 1974) and Miyata’s distances (Miyata et al. 1979). These measures are based on the volume and polarity of amino acids, as well as their chemical composition in the case of Grantham’s distance. For some applications, such measures are preferred over empirical distances, such as BLOSUM matrices (Henikoff and Henikoff 1992), for modelling substitutions in models that independently fit mutational parameters, as empirical matrices like BLOSUM include mutational biases already. Yet many other amino acid distance measures, and broader characteristics, that exclude mutational biases have been proposed, which warrant investigation for their relative utility in capturing amino acid exchangeability.

Among the most promising of these measures are those based on mutational scanning data: experiments where many amino acid replacements are tested in the same protein to evaluate their phenotype independently from mutation biases. Early forms of these experiments were used to develop the experimental exchangeability (EX) measure (Yampolsky and Stoltzfus 2005). In recent years, several deep mutational scanning experiments have been conducted, where every possible individual amino acid replacement is tested at each site of a protein, and across a wider set of proteins (Fowler and Fields 2014; Notin et al. 2023). Importantly, these amino acid substitutions are performed systematically, and are not based on random mutagenesis, meaning that this approach does not incorporate mutational biases. These new datasets open the possibility to update experimental exchangeability measures based on more extensive experimental data (Dunham and Beltrao 2021; Munro and Singh 2021).

Although amino acid distance measures warrant further investigation, these measures, by themselves, are ultimately simplistic. Indeed, some sites of a protein sequence are more critical than others for folding and function. As a result, several authors have argued that site-specific effects are also important to incorporate in molecular evolution models (Meyer and Wilke 2015; Hermans et al. 2025). Moreover, predicting the impacts of amino acid substitutions at particular sites in proteins (known as variant effect prediction) is an active area of research (Bromberg et al. 2024). There is naturally great interest in identifying which non-synonymous mutations are most likely to cause disease or modulate other phenotypes of interest. Methods developed for this purpose are generally of two classes: (1) sequence-based, which rely on deep alignments (Ng and Henikoff 2003) or protein language models (Rives et al. 2021; Elnaggar et al. 2022); (2) structure-based, which attempt to model protein folding and biophysics based on known or predicted structures (Gerasimavicius et al. 2025). In either case, benchmarking these approaches against simple amino acid distance measures that are not site-specific is important to assess their relative performance.

Here, we explore a range of existing amino acid distance measures. We also develop a new experimental exchangeability measure, DEX, which is the consensus of a separate measure that we computed based on recent deep mutational scanning data combined with the EX measure. We evaluated the relative utility of all measures for modelling codon substitution patterns and we also explored the utility of asymmetric exchangeabilities and the positions of residues within protein structure for predicting allele frequencies across highly divergent lineages. Our work provides a detailed comparison of the performance of existing measures and points a way forward for molecular evolution models that detect the strength and/or efficacy of purifying selection to incorporate more informative amino acid distance measures.

## Methods

### Existing amino acid dissimilarity measures

There are many approaches for capturing pairwise dissimilarity between amino acids, which have been aggregated into several databases and packages. Below, we briefly describe the dissimilarity measures we analyzed and where these values were acquired. Some of these measures represent similarity rather than dissimilarity: we took the complement of all measures as needed (after min-max scaling, see below) so that they all represent similarity or dissimilarity, depending on the analysis. For all asymmetric measures, before including them in codon substitution models, we took the mean of bidirectional distances between amino acids. Similarity measures were divided by the max similarity, to convert the max value to 1.0.

As mentioned above, the most well-known amino acid dissimilarity measures are Grantham’s distance (Grantham 1974) and Miyata’s distance (Miyata et al. 1979), which were partially motivated by two earlier measures. The first was Sneath’s D (Sneath 1966), which is based on 134 physicochemical characteristics of amino acids. The D index represents the percentage of these characteristics that are not shared between two amino acids. The second earlier measure is Epstein’s coefficient of difference (Epstein 1967), based on polarity and size. We parsed these four measures, as well as the EMPAR measure (Rao 1987), which is based on topological and physicochemical features, from the PARD python package (Lhotte and Taupin 2022). We manually parsed an additional dissimilarity measure, “Xia”, that captured which amino acids neighbour each other in sequences (Xia and Xie 2002). These authors reported their dissimilarity matrix as the average of their neighbour-based approach and Miyata’s distance, which we multiplied by two and from which we removed Miyata’s distance to back-calculate the neighbour score. A final existing matrix of pairwise amino acid (dis)similarities we parsed is the Conformational Similarity Weight (CSW) matrix (Kolaskar and Kulkarni-Kale 1992), which represents similarities based on pairwise comparisons of observed distributions of backbone dihedral angles (φ and ψ) for amino acids across the crystal structures of 102 proteins. The CSW matrix contains zero-valued entries, which would mean substitutions between certain amino acids are impossible, which would create issues in the codon substitution models described below. We replaced these values with a small positive value (0.01 x the matrix maximum, *i.e*., 0.1).

A different approach for computing amino acid dissimilarities is to summarize the impacts that substitutions have on protein function, based on datasets where substitutions in proteins have been experimentally investigated. This was the approach taken to develop the experimental exchangeability (EX) measure, that was based on experimental substitutions across 12 proteins (Yampolsky and Stoltzfus 2005). This measure, which we also acquired from PARD, was estimated based on fitting a power law model to observed impacts of each amino acid pair, on a variety of phenotypes. A similar measure, DeMaSk (Munro and Singh 2021), was recently developed based on 18 proteins where deep-mutational scanning has been performed, meaning nearly all possible amino acid substitutions were tested. These authors used an approach that involved transforming the experimentally obtained substitution impacts into rankings per site, to estimate the overall relative exchangeability per amino acid pair. The new dissimilarity measure we computed (DMS-EX) is closely related to these two prior measures but differs in the experimental datasets and transformation approach used, as described below.

Amino acid substitution matrices, such as BLOSUM62 (Henikoff and Henikoff 1992) and VTML200 (Müller et al. 2002), are well-known representations of pairwise substitution rates between amino acids, which share similarities with the above measures. The BLOSUM62 matrix is based on empirical differences between protein sequences with ≤62% identity, while VTML200 is a similar matrix but based on a maximum likelihood model of evolutionary divergence. Similar matrices have been estimated from empirical data, but which represent substitution rates rather than log-odds, the most well-known of which are JTT (Jones et al. 1992), WAG (Whelan and Goldman 2001), and LG (Le and Gascuel 2008). In all these matrices, amino acids with high substitution rates are generally more structurally similar and are therefore likely to have less impact on protein structure and function. However, such empirical matrices do not disentangle selective effects from mutational biases, which will also influence pairwise substitution rates between amino acids. Thus, we included BLOSUM62, VTML200, JTT, WAG, and LG as representative substitution matrices for comparison with other measures, but it would be inappropriate to use them to represent amino acid similarity in codon substitution models, where mutational biases are calculated separately. For BLOSUM62 and VTML200 we converted the log-odds scores to probabilities for our analyses.

Rather than estimating continuous differences, amino acid substitutions are often binned as conservative and radical, based on physicochemical groupings (Zuckerkandl and Pauling 1965; Hughes et al. 1990). We included these approaches in our molecular evolution models to assess whether continuous measures or binary classifications result in better model fits. We considered two distinct groupings (“Zhang RvC” and “Weber RvC”) based on polarity and volume (Zhang 2000; Weber and Whelan 2019), as well as groupings based on charge (Zhang 2000). To include these groupings in comparable analyses, we set amino acid pairs as having similarities of 99% or 1%, for conservative and radical substitution pairs, respectively.

The above measures are representative of how amino acid dissimilarity is typically captured in bioinformatic analyses. However, diverse amino acid characteristics, often based on dimension reduction approaches applied to many physicochemical properties, have been presented in the literature, which we also incorporated into our analyses. These approaches represent amino acid characteristics rather than proposed dissimilarity measures, as described below. We computed dissimilarity measures from each of these sets of characteristics by min-max scaling each individual variable (*i.e.,* scaled them to have a range between 0 and 1): 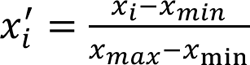

Where *x*_min_ and *x*_max_ are the minimum and maximum values for a variable across all amino acids, and *x*_i_ is the value for amino acid *i*. The Euclidean distance was then computed between amino acids based on all variables per measure: 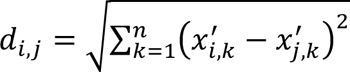, where *d*_i,j_is the distance between amino acids *i* and *j*, and *n* is the total number of variables. From the set of metrics aggregated in the Peptides R package v2.4.6 (Osorio et al. 2015), we parsed: (1) “Cruciani”: three scaled principal component scores (Cruciani et al. 2004); (2) “FASAGI”: six components representing diverse amino acid characteristics (Liang and Li 2007); (3) “Kidera”: ten orthogonal factors based on multivariate analysis (Kidera et al. 1985); (4) “zScales”: five factors representing physicochemical observations, including based on thin-layer chromatography and nuclear magnetic resonance data (Sandberg et al. 1998); (5) “VHSE”: 8 principal components based on hydrophobic, steric, and electronic properties (Mei et al. 2005). We similarly defined the “Atchley” measure based on five scaled factors previously published to summarize 54 amino acid features (Atchley et al. 2005).

Similar to the above methods which performed dimension reduction across amino acid characteristics, we repeated an analogous approach based on amino acid characteristics across different functional groupings in the AAontology database (Breimann et al. 2024). This database provides an ontology of amino acid scales, largely based on the AAindex database (Nakai et al. 1988; Tomii and Kanehisa 1996; Kawashima and Kanehisa 2000; Kawashima et al. 2008), which itself parsed amino acid characteristics from over 100 articles. The AAontology (“AA-Ont.”) database includes 586 amino acid scales split across eight categories: composition (“Comp.”), conformation (“Conf.”), energy, others, polarity (“Pol.”), shape, structure/activity (“Struc.”), and volume (“Vol.”). We computed principal coordinates analyses across all scales per category and computed a single dissimilarity measure per category by performing a similar calculation as above. Rather than min-max scaling, we retained the original values and instead weighted the difference in each principal component by the variance explained: 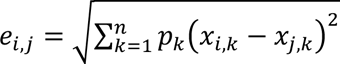, where *e*_i,j_ is the distance between amino acids *i* and *j*, *n* is the total number of principal components included, and *p*_k_ is the proportion of variance explained by component *k*. We used this same approach to generate a single dissimilarity measure (“Venkatarajan”) from another analysis that provided five factors (and corresponding eigenvalues) representing 237 physicochemical properties (Venkatarajan and Braun 2001).

To compare all these measures visually, we computed the mean exchangeabilities per amino acid for each measure. These exchangeability values were conceptually based on Graur’s stability index (Graur 1985), but based on amino acid similarities rather than distances. Specifically, we computed the mean amino acid similarity based on every possible single non-synonymous mutation in a codon. We visualized these results in a clustered heatmap based on taking the standard score of the exchangeabilities per measure and then computing Euclidean distances for both amino acids and measures (*i.e.,* for rows and columns separately) and then performing hierarchical clustering with the complete method. We also ran Principal Component Analyses (PCAs) on the exchangeabilities scaled per measure to visualize the relative similarity of measures. We built one PCA based on all measures and one PCA with these major outlier measures removed, for better visualization: CSW, EMPAR, Comp. (AA-Ont.), Conf. (AA-Ont.), RvC (charge), RvC (Zhang), and Venkatarajan.

### Re-calculating classic distance measures

In addition to the standard Grantham’s distance values, we re-calculated Grantham’s distance based on the composition, polarity, and molecular volume values presented in the original publication (Grantham 1974). We did so because we noticed rounding errors in the original publication. We calculated the raw Grantham’s distance between amino acids *i* and *j* as: 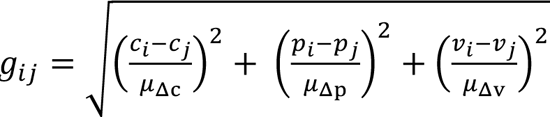, where *c*, *p*, and *v* represent composition, polarity, and volume, and *μ*_Δc_, *μ*_Δp_, and *μ*_Δv_ represent the mean absolute difference between all amino acid pairs based on each variable alone (*μ*_Δc_ = 0.7393684, *μ*_Δp_=3.134211, and *μ*_Δv_= 50.06053). We then calculated the scaled Grantham’s distances, *G*_i j)_, which are scaled so that the mean distance across all pairs is 100, following from the original calculation: *G*_i j)_ = *g*_i j)_ × (100/*μ*_6_), where *μ*_6_is the mean raw Grantham’s distance between all amino acid pairs (*μ*_6_=1.968261). These final scaled Grantham’s distances are similar to the originally published distances (mean difference in values: 0.03), but there are some noteworthy differences, including a maximum absolute difference of 9.83 units between the original and re-calculated distances (for the distance between aspartic acid and tryptophan, which is under-estimated in the original table).

Grantham’s distance is the most commonly used, general-purpose, amino acid distance measure, *e.g*., (Graur 1985; Chowell et al. 2019). However, substantial work has been conducted to further refine estimates of amino acid properties since it was originally published. Accordingly, we also re-calculated Grantham’s distances with updated volume and polarity estimates. We did not adjust the composition estimates as these were specifically defined for the original measure. To this end, we used more recently computed amino acid volumes that have been shown to have lower variance compared to other approaches (Tsai and Gerstein 2002). For a measure of polarity we followed the same approach of scaling and then averaging the polar requirement (Woese et al. 1966) and a measure of hydrophobicity per amino acid. We used an estimated measure of polar requirement based on molecular dynamics simulations, rather than the original values, as the former avoids systematic experimental error (Mathew and Luthey-Schulten 2008). In addition, rather than the hydrophobicity measure that was originally used (Aboderin 1971), we used a measure that has been shown to perform best (out of 98 hydrophobicity measures) for distinguishing peptides based on hydrophobicity features (Naderi-Manesh et al. 2001; Simm et al. 2016). We then multiplied the hydrophobicity metric by −1 (to make the two variables positively correlated) and performed min-max normalization of the two measures across the amino acids (so that they each range from 0 to 1). We then computed the mean of these scaled values to represent each amino acid’s polarity.

We used these same steps to re-calculate Miyata’s distances based on these updated hydrophobicity and volume measurements. Raw Miyata’s distances were calculated following from the original approach (Miyata et al. 1979): 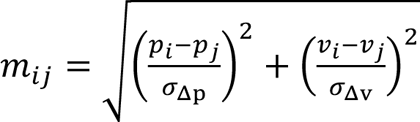, where *σ*_Δp_ and *σ*_Δv_ are the standard deviations in pairwise differences in polarity and volume between all amino acids. We conducted a supplementary comparison of these varied computations of Grantham’s and Miyata’s distances, to ensure that our conclusions regarding these well-known measures did not depend on minor differences in how they are computed.

### Computing a new experimental exchangeability measure

We downloaded all 217 deep mutational scanning datasets from the ProteinGym database v1.1 (Notin et al. 2023). We then strictly filtered these datasets down to a smaller high-quality and independent set. This involved first retaining only datasets with at least 95% of sites tested (minimum of 20) and a mean of at least 15 (out of 19) amino acids tested per site. We then subset the target sequences that were used for the experiments to be only residues where all 19 replacement amino acids were experimentally tested (with multi-substitutions excluded). We also dereplicated datasets with similar proteins by clustering all target protein sequences with CD-HIT v4.8.1 (Fu et al. 2012) at 70% identity and keeping only the representative sequence for each cluster. We also determined how each score was transformed based on the original published articles and excluded scores with ambiguous or unclear transformations. At this point, 79 datasets remained for which we could unambiguously determine what the score represented in terms of the transformations employed. These included 60 datasets from the same study (Tsuboyama et al. 2023). We decided to keep only one dataset per independent publication to help avoid biases and to include substitution impacts across more varied proteins and experimental protocols, which resulted in a final set of 18 deep mutational scanning datasets (**Supplementary Table 1**) (Melnikov et al. 2014; Klesmith et al. 2015; Majithia et al. 2016; Weile et al. 2017; Wrenbeck et al. 2017; Lee et al. 2018; Mighell et al. 2018; Soh et al. 2019; Chen et al. 2020; Jia et al. 2021; Chen et al. 2023; MacRae et al. 2023; Suphatrakul et al. 2023; Tsuboyama et al. 2023; Weeks and Ostermeier 2023; Clausen et al. 2024; Estevam et al. 2024; Vanella et al. 2024). For datasets representing raw scores we set a minimum score of 0.01. We then transformed all the scores across these datasets to a natural log scale.

Following the approach developed to generate the EX measure (Yampolsky and Stoltzfus 2005), we treated substitution scores separately depending on whether they were derived from buried or exposed regions of the proteins. To identify these regions, we analyzed all AlphaFold2 structures (Akdel et al. 2022) associated with the target protein sequences. We first ran mkdssp v3.0.0 (Touw et al. 2015) to identify secondary structure features, including solvent accessibility (Kabsch and Sander 1983). We then calculated the relative solvent accessibility per residue using previously determined maximum allowed solvent accessibilities per amino acid and a Python script to conduct this calculation (Tien et al. 2013). We defined buried sites as those with a relative solvent accessibility 0.2, with all others defined as exposed. We used the same workflow to compute the relative accessibility of all amino acids across the AlphaFold-predicted structures across the entire *Escherichia coli*, human and yeast proteomes (v4) (Akdel et al. 2022). We then calculated the mean proportion at which each amino acid is found within buried and exposed sites, for each species separately. We then took the mean of these three mean values per amino acid to get an overall estimate of the background proportion of cases that each amino acid is found at buried vs. exposed sites.

To calculate our custom deep-mutational scanning exchangeability score, we considered all residues in the 18 target proteins with at least 15 amino acid substitutions tested. For each amino acid substitution type, and within each tested protein and site-type (buried or exposed), we computed robust Z-scores. Specifically, for all exposed (*e*) residues, we computed the robust Z-score (*r*) for a given substitution (*j* → *k*) at site *i* in protein *p* as: 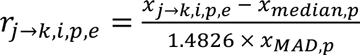. Where x_median,p_ is the median score for all substitutions in this protein and *x*_MAD.p_ is the median absolute deviation of all scores. The analogous approach was used at buried sites as well. Importantly, the median and median absolute deviation are computed at all residues in the protein, both buried and exposed. The median absolute deviation is multiplied by 1.4826 to approximate the standard deviation when the underlying data is normally distributed (Rousseeuw and Leroy 1987). We then computed the median robust Z-score per amino acid substitution type, per protein, and for buried and exposed sites separately, and then computed the median of these values across all genes, meaning that for each amino acid substitution type there is a score for the buried and exposed sites separately (*r*_j→k,C_ and *r*_j→k,9_). Finally, we computed a weighted score per substitution type: *s*_j→k_ = *r*_j→k,C_ × *b*_j,b_ + *r*_j→k,e_ × *b*_j,b_, where *b*_j,b_ and *b*_j,e_ are the background frequencies of the amino acid *j* across the background proteomes at buried and exposed sites, respectively. We refer to this custom measure as DMS-EX.

We also computed consensus measures based on subgroups of measures using the DISTATIS approach (Abdi et al. 2012) with the DistatisR package v1.1.1. DISTATIS yields compromise factor scores (analogous to PCA scores) based on multiple distance matrices. In this case, the factor scores capture where amino acids are positioned in the compromise space, weighted by the eigenvalues. We retained all factors with positive eigenvalues and computed a consensus measure for each grouping of measures by computing the Euclidean distances between pairwise amino acids based on these factors.

### Fitting codon substitution models to diverse alignments

We fitted codon substitution models on separate gene alignments across six *Drosophila*, 190 mammal, and 71 *Streptococcus* species, to ensure that a diverse set of independent lineages was evaluated. *Drosophila* and mammal alignments were chosen given that these lineages are well-studied and have resources available. We acquired the available alignments and species trees for these lineages from flyDIVaS v1.2 (Stanley and Kulathinal 2016), for the six species in the “melanogaster species group”, and from orthoMaM v12 for the mammalian data (Allio et al. 2024).

We generated custom alignments of the *Streptococcus* species. We selected this genus as a representative of prokaryotes which has high species diversity among sequenced isolates. We downloaded 3,270 genomes from GenBank and then identified 71 different *Streptococcus* species defined at a 95% average nucleotide identity using skani v0.2.2 (Shaw and Yu 2023). We selected one genome for each of these species clusters and ran Bakta v1.8.1 (Schwengers et al. 2021) followed by PanTA v1.01 (Le et al. 2024) to produce a pangenome breakdown across the genus, and identified all gene families found in at least 68/71 genomes. We then ran MACSE2 (Ranwez et al. 2018), specifically the OMM_MACSE v12.01 workflow, to build codon alignments for each of these gene families across these genomes and built trees for each gene family with fasttree v2.1.11 using default settings (Price et al. 2010).

For the three lineages, we decided to concatenate genes together to ensure sufficient data for model fitting because running codon models on every individual gene would have limited numbers of substitutions for fitting the models robustly. Before doing so we estimated the numbers of synonymous and non-synonymous substitutions across all gene alignments across all three lineages using Fitch parsimony (Fitch 1971). We retained only alignments with a length between 200-1000 codons, inclusively, and with at least 20 synonymous and 20 non-synonymous substitutions. We then identified 20 non-overlapping, random sets of 12 genes for each lineage and concatenated these 12 gene alignments together for each random set. Only genomes with all 12 genes present were included within each set. We generated new trees per alignment set with fasttree for the mammalian and *Streptococcus* concatenated alignments.

To fit the substitution models, we ran CODEML from the PAML package v4.10.7 (Yang 2007), as wrapped in the Biopython Python package v1.85 (Cock et al. 2009). We ran CODEML on all alignments (described below) with both the M0 model (standard d_N_/d_S_) and with an amino acid distance matrix corresponding to the measures described above, which was specified with the ‘aaDist’ option (Goldman and Yang 1994). We compared Bayesian Information Criterion (BIC) values across the fitted models ranked from lowest (best) to highest, where dataset size was taken to be the number of codons used per model. We ran the first 10 sets of alignments on all individual measures and used these to identify the best-performing measures, which we then used as the basis for the consensus amino acid measures produced with DISTATIS, as described above. We evaluated these consensus measures on the remaining 10 sets of alignments.

### Segregating non-synonymous variants across humans and *E. coli*

We downloaded segregating non-synonymous variants in 4,335 proteins across 2,661 wild-type *E. coli* strains (Catoiu et al. 2023) that are part of the Alleleome dataset (https://github.com/EdwardCatoiu/Alleleome/). We only considered sites that contained unambiguous amino acid residues for at least 2,000 strains. We also ignored substitutions at the first residue per protein, limited to substitutions caused by a single mutation, and retained only the most frequent substitution per position. Singleton variants (those found in a single strain) were also excluded due to the likelihood of these being identified by sequencing error. Only proteins with clear AlphaFold structural predictions (see below) were retained for analyses. We polarized all mutations (i.e., we identified which allele is the ancestor vs. derived, to produce an unfolded allele frequency spectrum) by identifying the amino acids at each orthologous residue present in close relatives of *E. coli*. The relatives we focused on were *Escherichia fergusonii* (genome: GCF_000026225.1) and *Salmonella enterica* (genome: GCF_000006945.2). We identified bidirectional best-hits with Protein-Protein BLAST v2.17.0+ (Altschul et al. 1990) between the proteomes of these species and the consensus proteins from the Alleleome database. We identified 1:1 orthologs as those that bidirectionally best matched each other based on: E-value ≤10^-10^, coverage of at least 90% (for subject and query), and bit-score ≥10% higher than the next best query match. We aligned all orthologs using MAFFT v7.525 (Katoh and Standley 2013) with local-pair mode and a maximum of 1,000 iterations. We retained only amino acid residues where both outgroup species were the same for downstream analyses, and which matched either the minor or major allele in *E. coli*, to unambiguously infer the ancestral state.

We similarly downloaded and parsed segregating non-synonymous mutations across human genotypes. We downloaded all exonic VCFs (*i.e.,* for all chromosomes) from the gnomAD variation database (v4.1) (Karczewski et al. 2020). These variant calls include annotations of non-synonymous calls across all proteins. We downloaded protein and transcript sequences from GENCODE v45 (Frankish et al. 2023), and identified the proteins for which RaSP predictions were available. We then used a custom Python script to parse out non-synonymous variants across all chromosomes, including information on the amino acid residue in the protein, the amino acid change, and the frequency. The remaining steps were the same as the *E. coli* analysis above, where the GENCODE protein (and codon) sequences were used as the references for predicting site-specific alleles. We did not use a minimum allele frequency threshold for variants in the gnomAD variation database, given the high-quality nature of the gnomAD database, but required a minimum of 1,400,000 individuals per site. We polarized all mutations by identifying the amino acids present at orthologous sites in the chimpanzee (Pan_tro_3.0), gorilla (gorGor4), and the Sumatran orangutan (PABv2) reference genomes. We identified all one-to-one orthologs using ENSEMBL BioMart (Kinsella et al. 2011) and then aligned these proteins with the focal human proteins using MAFFT as above. We restricted our analysis to sites where all three outgroup species agreed on the ancestral amino acid residue, and where this matched either the human minor or major allele.

We fit linear models to two allele frequency-based dependent variables: mean allele frequency and the proportion of variants classified as rare (<0.1% and <0.0001% for *E. coli* and human, respectively). Each observation included each possible (asymmetric) amino acid substitution (represented by at least 10 substitutions in both buried and exposed sites separately), for which we computed the two allele-frequency dependent variables. To perform a basic correction for mutational biases, we tracked whether the underlying codon substitutions were driven by transversions or transitions. In humans, we also tracked whether transitions occurred at CpG sites. We then defined categorical variables representing the most common type of mutation driving each amino acid substitution. In *E. coli* this was represented simply as transitions being present in >50% of cases or not. In humans we classified mutations as transversions, non-CpG transitions, and CpG transitions, using this same majority rule (and where there was a clear majority case for most substitutions). All continuous variables were transformed to normal distributions using the orderNorm transformation (Peterson 2021), using the R package bestNormalize v1.9.1, to allow a standard linear modelling approach to be used.

### Running site-specific effect predictions

To provide context for how informative amino acid dissimilarity measures perform at predicting deleterious amino acid substitutions compared to state-of-the-art methods, we also analyzed the outputs from three deep learning approaches. These include RaSP (Blaabjerg et al. 2023) and ThermoMPNN (Dieckhaus et al. 2024) for protein structure-based predictions and VespaG (Marquet et al. 2024) as a method based on embeddings from protein language models. These approaches are representative of the two main approaches used for predicting substitution effects: structure and sequence-based methods. RaSP is computationally intensive to run and so we opted to use pre-existing predictions based on predicted structures across the human proteome (https://doi.org/10.17894/ucph.7f82bee1-b6ed-4660-8616-96482372e736). Unfortunately, no such structural predictions were available for *E. coli* to our knowledge. We ran ThermoMPNN (v1.0.0) on AlphaFold-predicted structures (the same set analyzed to produce our primary metric). We ran VespaG (commit: 660b17c6964eb6db8e3f3bee2b8bbd3f9f574d23) with the ESM-2 protein language model embeddings (Lin et al. 2023).

### Visualization and working environments

Commands were run using the R (v4.2.2) and Python (v3.10) programming languages. Plots were generated using the R packages ggplot2 v3.5.1 (Wickham 2016) and ComplexHeatmap v2.20.0 (Gu 2022). We installed command-line tools from the Bioconda project (Grüning et al. 2018), when possible, and parallelized commands using GNU Parallel version 20240222 (Tange 2011). We used Claude AI to help check for errors across our scripts and manuscript.

## Results

### Exploring existing amino acid distance metrics and introducing a new measure

Many different measures capturing amino acid distances have previously been proposed (**Table 1**; **Supplementary Table 2**). These include binary measures of radical vs. conservative substitutions, as well as many scales based on physicochemical properties. A more recent set of approaches is based on experimentally observed impacts of amino acid substitutions in proteins (Yampolsky and Stoltzfus 2005; Munro and Singh 2021).

**Table 1.** Highlighted pairwise amino acid distance measures analyzed in this study.

| Group | Approach | Description |
| --- | --- | --- |
| Radical vs. conservative (RvC) | RvC groupings based on polarity and volume (Zhang 2000) | Classifying substitutions as radical or conservative based on whether amino acids have highly different polarity and volume values (radical substitution) or not (conservative substitution). |
| Physicochemical/ structural | Grantham’s distance (Grantham 1974) | Distance based on values of composition, polarity, and volume for each amino acid. |
| Substitution-based (empirical) | BLOSUM62 (Henikoff and Henikoff 1992) | Based on observed substitutions across protein alignments where sequences are |
|  |  | ≤62% identity. Confounded with nucleotide mutational biases. |
| Experimental | Experimental exchangeability (EX) (Yampolsky and Stoltzfus 2005) | Based on a comparison of 9,671 introduced amino acid substitutions across 12 proteins. |
| Experimental | DMS-EX (this study) | New measure of experimental exchangeability that we computed based on deep-mutational scanning datasets. |
| Experimental | DeMaSk (Munro and Singh 2021) | Similar to DMS-EX and EX but based on a slightly different set of datasets and based on rank-based transformations. |
| Experimental | DEX: DISTATIS-based consensus of experimental exchangeability (this study) | New measure that we computed as the consensus of EX and DMS-EX, using DISTATIS. It is the top-performing measure across all the matrices that we evaluated. |

Partially motivated by these approaches, we computed a new measure, DMS-EX, which is based on publicly available deep mutational scanning datasets (Notin et al. 2023). DMS-EX differs from past approaches in terms of the exact datasets included and especially because it incorporates the magnitude of impact differences, which are all on the same scale across datasets, rather than just the relative rankings (see Methods).

To visualize the similarity across existing measures, in addition to our novel measure, DMS-EX, we performed a comparison of 30 differing measures (see Methods for a detailed description of these measures). For each measure we computed the exchangeability per amino acid, which represents the mean similarity to all other amino acids that can be reached by one non-synonymous mutation. We then clustered these exchangeability values to visualize variation across measures (**Figure 1a**). At a broad level, the measures are consistent with one another in terms of dividing amino acids like tryptophan and arginine (mean standard scores < −1.1) with low exchangeability from others with higher exchangeability, such as glutamine and methionine (mean standard scores >0.65). As expected, the measures are correlated to one another on average (median Spearman’s ρ=0.45; standard deviation: 0.27), and we observed groupings between similar measures, such as DMS-EX with EX and DeMaSk, which are all based on experimental data. However, certain measures are major outliers driving the variation (see Methods; **Supplementary Figure 1**). Excluding these outliers, we visualized the relative similarity in these measures based on the first two principal components (**Figure 1b**). A single component explains a substantial amount of the variance (64.7%), which is consistent with clustering driven by two main types of approaches: those based on physicochemical measures (*e.g*., RvC and Grantham) and those based on empirical substitutions (*e.g*., BLOSUM62 or EX). We observed that a small number of amino acids display an inflated impact on the variance across measures, particularly tryptophan and arginine, rather than an overall shift in exchangeability across all amino acids (**Figure 1c**).

**Figure 1:**
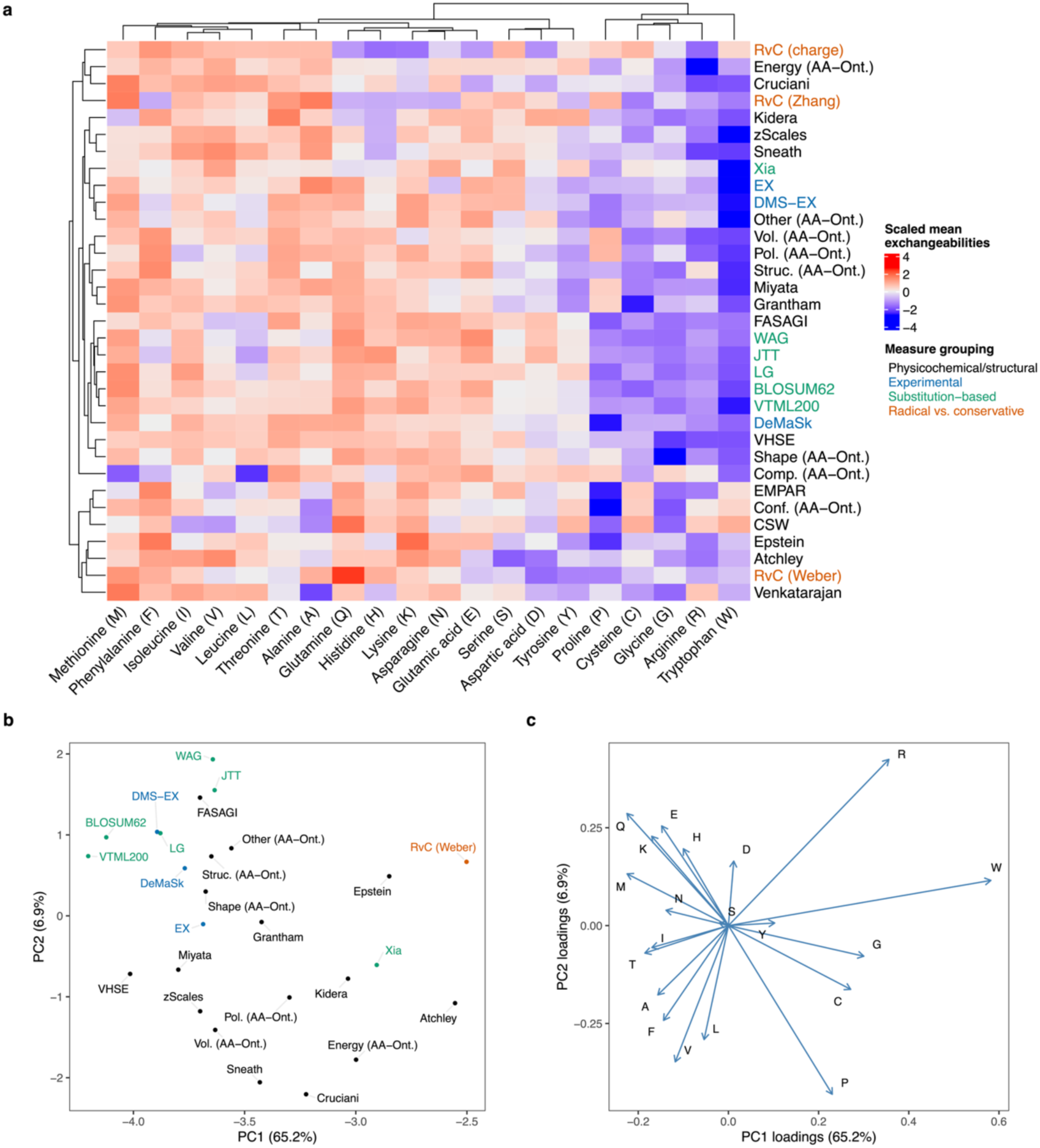
Comparison of amino acid (dis)similarity measures. (a) Mean exchangeabilities per measure, averaged across all codons for each amino acid. A codon’s mean exchangeability was calculated as the mean amino acid similarity between all amino acids that could arise from a single non-synonymous mutation to the codon. Rows represent the different metrics analysed (which were all converted to similarities, see the main text for description). Scaled values were hierarchically clustered based on Euclidean distances using the complete method, for both the rows and columns separately. (b) First two principal components (PCs) of Principal Components Analysis (PCA) of all dis(similarity) measures based on Euclidean distances. Outlier measures (CSW, EMPAR, Comp. (AA-Ont.), Conf. (AA-Ont.), RvC (charge), RvC (Zhang), and Venkatarajan) were excluded from this PCA for better visualization (**Supplementary** Figure 1). (c) Same PCA as in panel b but displaying the loadings that each amino acid contributes to the separation. Percentage of variance explained by each PC is indicated in parentheses.

### Fitting models of protein-coding gene evolution with amino acid distance measures

To assess how well these amino acid distance measures explain actual non-synonymous substitutions patterns, we applied a previously developed codon substitution model that incorporates an amino acid distance matrix (Goldman and Yang 1994). We applied this model to diverse alignments representing 71 *Streptococcus* species, six *Drosophila* species, and 190 mammalian species. We constructed 10 alignments of 12 concatenated genes randomly sampled (non-overlapping) for each of these independent lineages and applied the codon substitution model for each amino acid distance matrix to each separate alignment.

Based on the rankings of model performances across these alignments (**Figure 2**), all models that incorporated amino acid distances performed better, as expected, than the standard codon substitution model without a distance matrix (i.e., where all non-synonymous substitutions are equally weighted). Our custom measure, DMS-EX (mean rank: 1.17) and the EX measure (mean rank: 1.97) performed best on average across all three lineages. The difference in Bayesian Information Criterion (BIC) values in models based on these two measures was significantly different from 0 (DMS-EX resulted in BIC values 715.3 lower values on average; Wilcoxon test, *P*<0.001), indicating that DMS-EX fits significantly better overall. In addition, because Grantham’s and Miyata’s distances are so well known in molecular evolution, we updated these measures since more accurate estimates of the volume, polarity and hydrophobicity of amino acids are now available (see Methods). Interestingly, these updated values provided no clear improvement over the original measures (**Supplementary Figure 2**).

**Figure 2:**
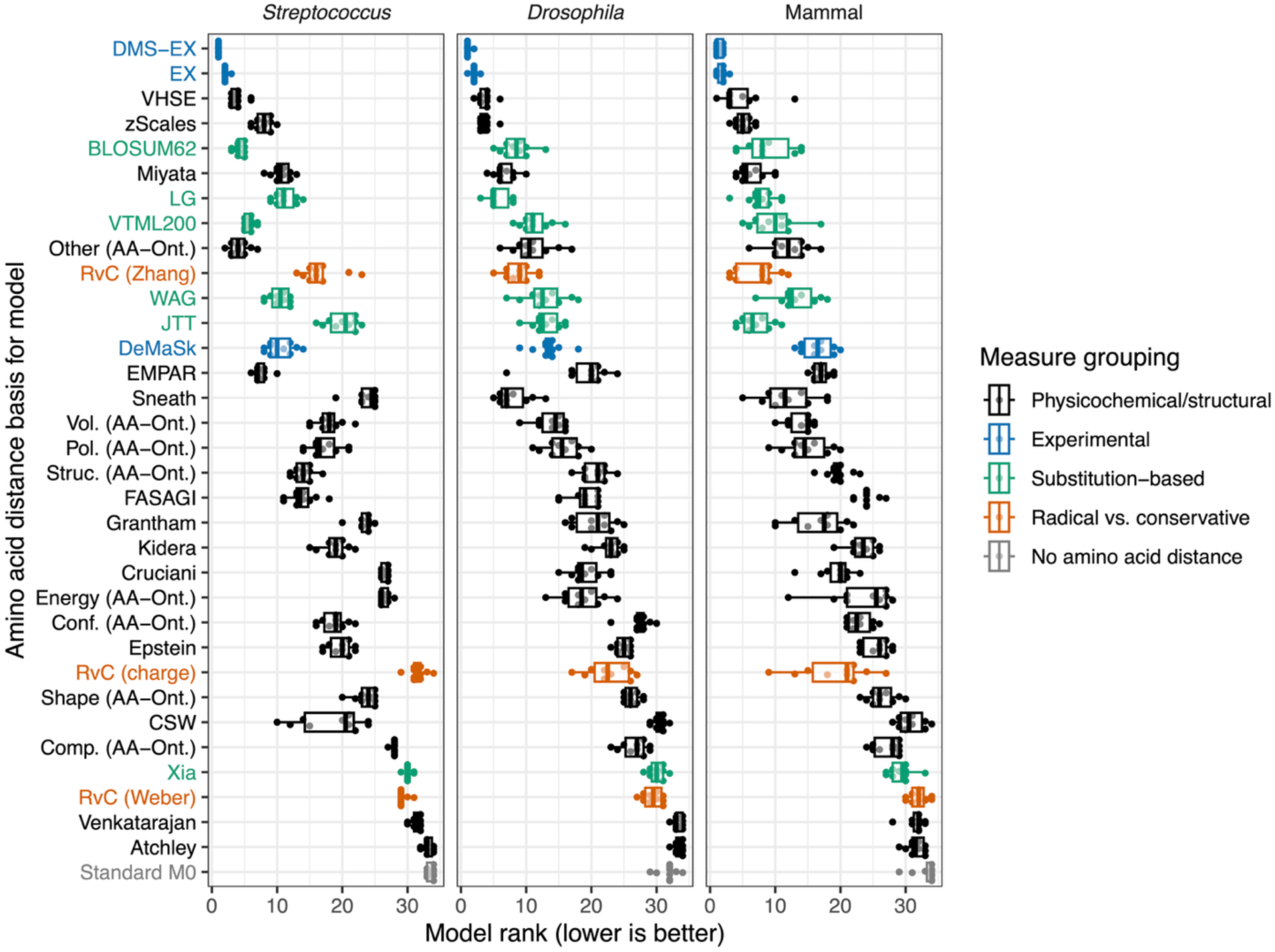
Comparison of model rankings, based on Bayesian Information Criterion values, for models applied to 10 different protein-coding gene alignments (with each alignment containing 12 randomly selected concatenated genes) across three highly diverged lineages using different amino acid similarity metrics.

We reasoned that a consensus measure based on several top-performing measures could perform best for modelling non-synonymous substitution patterns. To this end, we produced numerous consensus measures with the DISTATIS approach (see Methods) by generating different combinations of the top six performing measures (excluding BLOSUM62 since it incorporates mutational biases). We also evaluated consensus measures incorporating DeMaSk, because this measure was constructed using a similar approach to the one we used to generate DMS-EX. We evaluated these consensus metrics on a new set of 10 alignments for each of the three lineages (**Supplementary Figure 3**). We found that the separation between the top measures was less clear, but the consensus measure that combined our DMS-EX approach and EX was ranked best on average (mean rank: 2.3). We named this new consensus measure “DEX”, standing for “DISTATIS-based consensus of experimental exchangeability”. More generally, we found that DEX fits best across all alignments compared to the individual measures based on model rank (**Figure 3**). In addition, the difference in Bayesian Information Criterion values for models fit on DEX vs. DMS-EX (which is the next best-performing measure) were significantly greater than 0 (median difference: 161.45; Wilcoxon test *P*<0.001).

**Figure 3:**
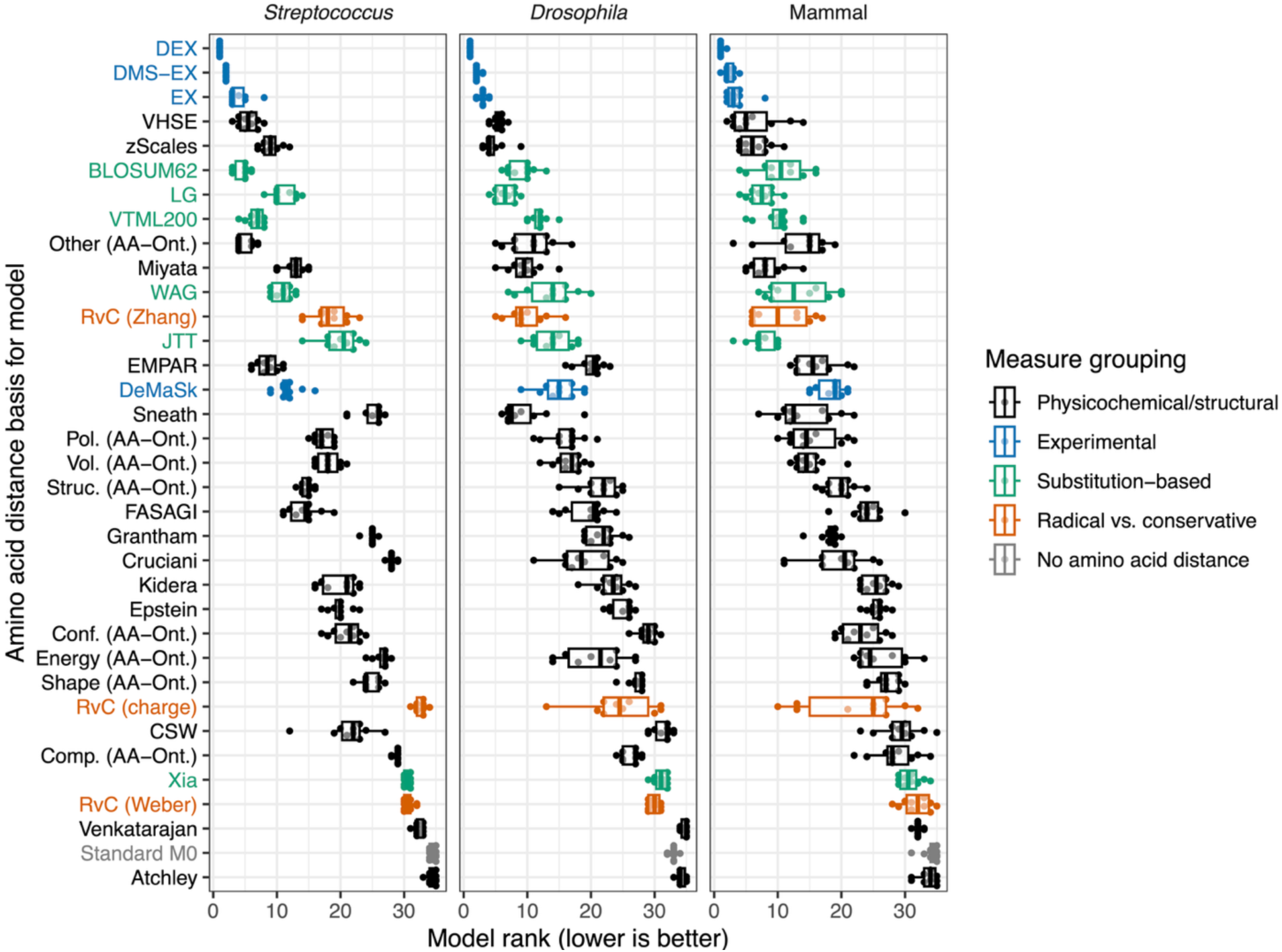
Comparison of model rankings, based on Bayesian Information Criterion values, for models applied to 10 different protein-coding alignments across three highly diverged lineages. Each model includes a different amino acid distance measure to represent pairwise dissimilarity. This result corresponds to an independent set of codon alignments compared to the earlier analysis used to identify the top measures to test in new combinations.

### Symmetric exchangeabilities are most associated with observed segregating amino acid polymorphism frequencies

The above analyses focused on symmetric matrices, where the exchangeability of amino acids such as K→L is the same as L→K. This is a requirement of standard codon substitution models, which require symmetric exchangeability rates in general. However, one benefit of using empirical DMS data to infer exchangeabilities is that it provides asymmetric information. In addition, we inferred the exchangeability of amino acids at buried and exposed sites separately, by first classifying residues into site classes by their relative solvent accessibility (RSA), before integrating them into the focal symmetric DMS-EX measure we analyzed above (see methods). Our results show that the asymmetric DMS-EX measures by burial site capture more nuanced patterns (**Figure 4**). In particular, we found a clear difference in exchangeability scores at buried and exposed sites. The asymmetric DMS-EX score for substitutions at exposed sites is on average 1.96-fold (median: 1.55-fold) higher than the corresponding substitution at buried sites (Paired Wilcoxon Test *P*<0.001). This is an expected result, as buried residues are known to evolve under stronger overall purifying selection (Goldman et al. 1998; Franzosa and Xia 2009). There are also certain amino acid pairs that show discordant asymmetry patterns between buried and exposed sites. For instance, C and V have relatively symmetrical exchangeability at buried sites, but C→V is substantially less exchangeable at exposed sites specifically relative to V→C (**Figure 4**).

**Figure 4:**
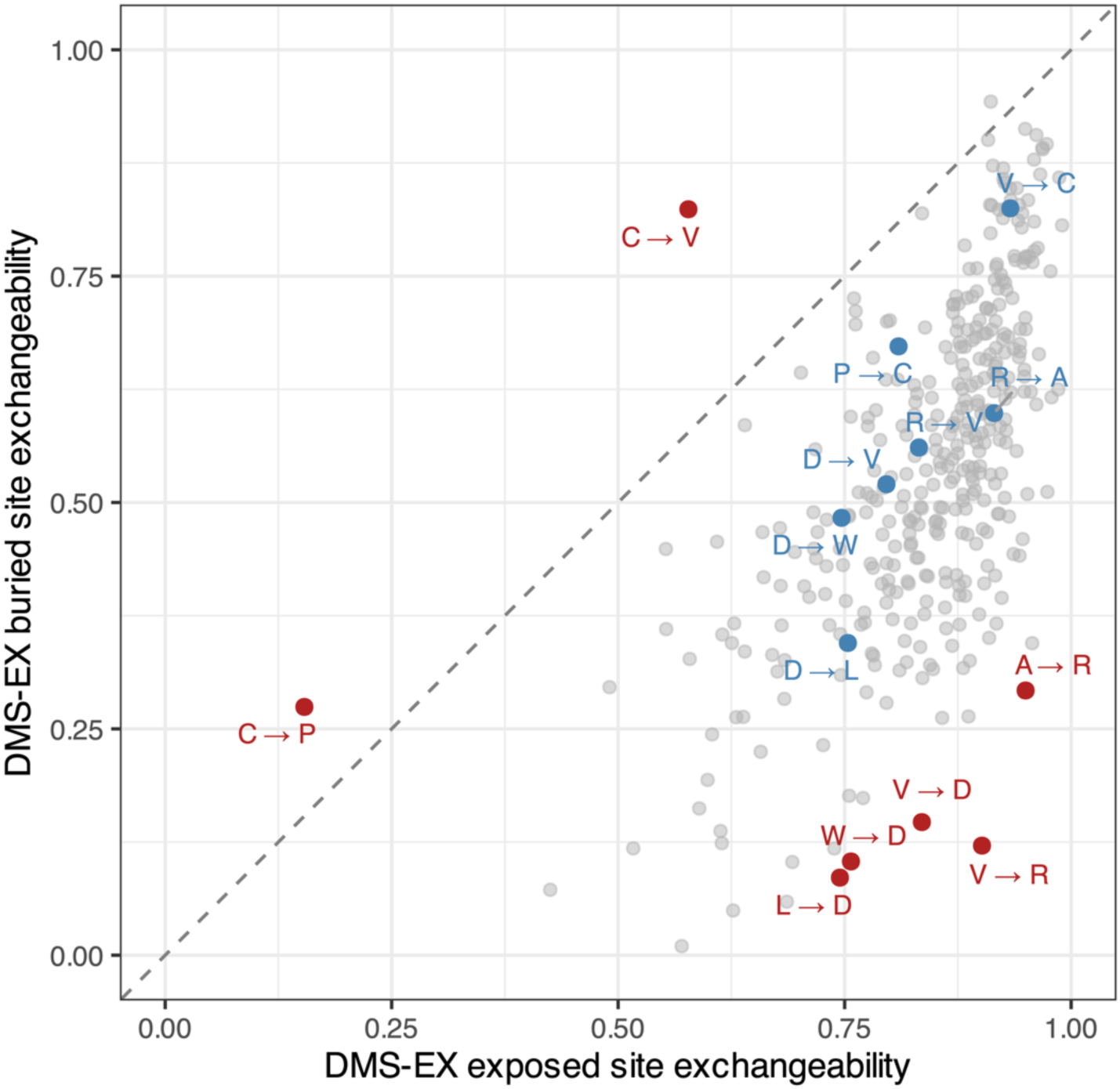
DMS-EX exchangeabilities for all asymmetric pairs of amino acids, split by the exchangeability at buried and exposed sites. A weighted average of these scores was used for the focal DMS-EX score, but the exchangeabilities at buried and exposed sites can also be analyzed independently. Highlighted outliers that differ in exchangeability by burial status are indicated in red, with the corresponding substitutions in the opposite direction indicated in blue. The dotted line indicates the 1:1 line.

We investigated whether asymmetric DMS-EX exchangeabilities provide improved insight over the corresponding symmetric measure. To do so, we compared whether symmetric or asymmetric versions of our focal measures explained more variance in segregating amino acid substitutions’ allele frequencies. Due to the action of purifying selection, deleterious mutations are less likely to rise to high frequencies in populations. We hypothesized that asymmetric exchangeability measures would account for more variance in observed allele frequencies, due to the increased information content of these approaches. We focused on two very large datasets of highly divergent lineages to once again ensure that our results were generalizable: *Escherichia coli* and humans. The final set of *E. coli* variants included 38,802 segregating replacement mutations across 1,802 proteins and a minimum of 2,000 wild-type strains. We computed the mean allele frequency for each pair of amino acids per substitution. We performed a similar analysis on human population data, which included 4,224,357 non-synonymous variants across 11,795 protein coding genes obtained from the gnomAD human variation database. Using these datasets, we fit linear models to predict mean allele frequency using RSA group (buried or exposed), mutation type, and exchangeability score. Each observation was thus the mean allele frequency of each asymmetric amino acid substitution at buried or exposed sites. All continuous variables were transformed with the orderNorm transformation to enforce normality, and thus these analyses focus on relative rankings rather than absolute magnitude and original units.

We considered two asymmetric versions of the DMS-EX exchangeability matrix. The first (DMS-EX weighted) is the asymmetric matrix but as a weighted average over buried and exposed sites. In contrast, the second (DMS-EX by RSA group) retained separate exchangeability values for buried and exposed sites. This partitioning allowed us to investigate whether asymmetry in general, or asymmetry conditioned on buried or exposed sites specifically, results in better model fitting. Surprisingly, in both species, symmetric exchangeability matrices (either DMS-EX or DEX) resulted in the best fitting model (**Figure 5**), albeit with a small difference in effect size in the human data. We found the same overall result when predicting the proportion of rare alleles as the predictor rather than mean allele frequency (**Supplementary Figure 4**). In addition, although we focused on our focal metrics (DMS-EX variants and DEX), the general pattern of symmetric metrics better explaining the variance in allele frequency was a robust signal overall (**Supplementary Figures 5 and 6**).

**Figure 5:**
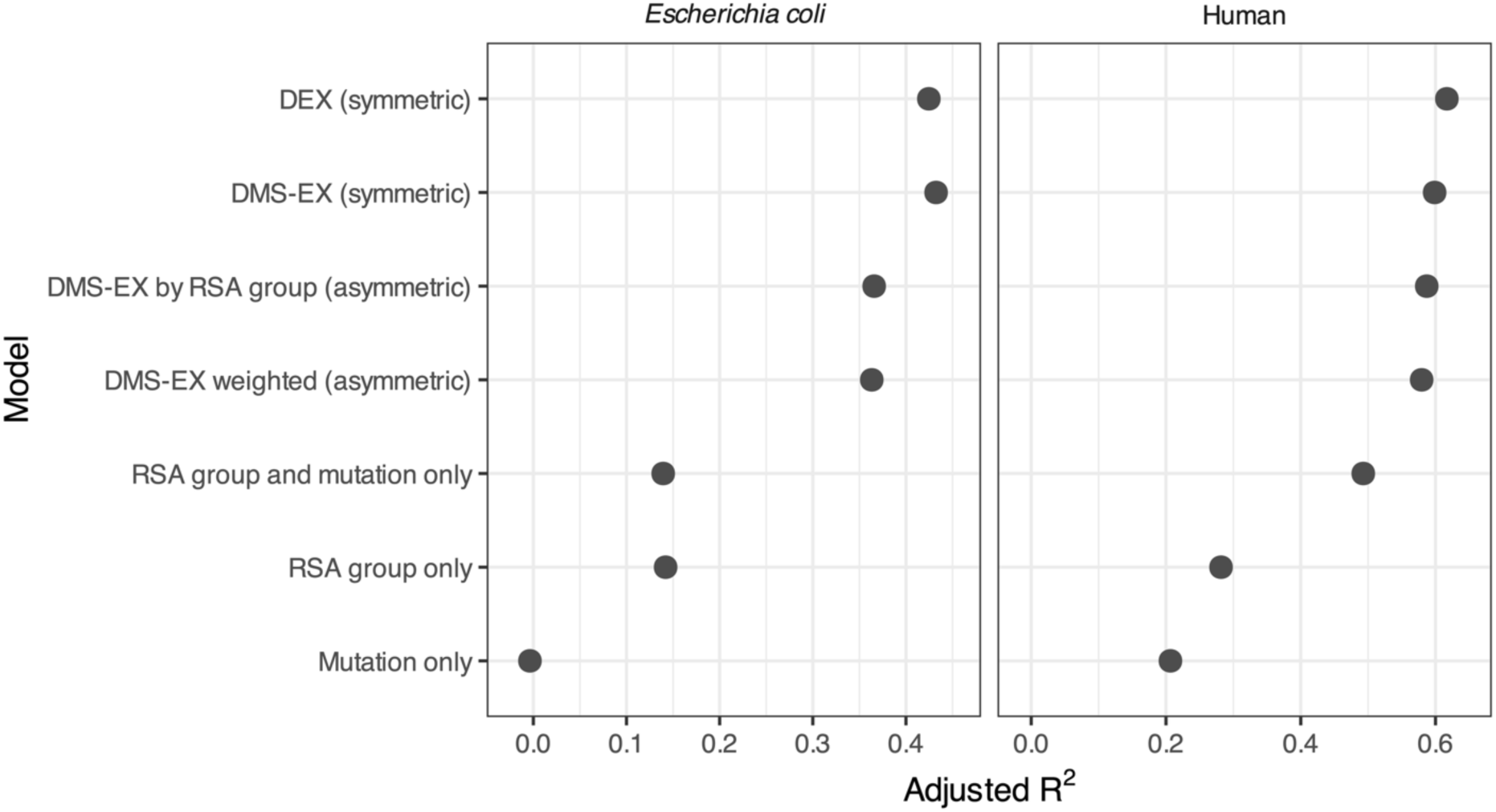
Adjusted R^2^ values for linear models predicting mean allele frequency for segregating amino acid replacements across *E. coli* and human populations. See main-text for a description of the asymmetric variables. The symmetric DEX and DMS-EX measures match the focal metrics used in the earlier codon substitution models. The models with RSA and/or mutational coefficients only are included to provide baselines over which exchangeability matrices provide an improvement. Note the differing relative importance of controlling for transition vs. transversion mutations (the mutational model in *E. coli*, whereas in the human models we also partitioned transitions by CpG sites) in each species. RSA: Relative Solvent Accessibility.

## Discussion

We developed a new consensus-based experimental exchangeability measure, DEX, and showed that this measure best fits real codon substitution patterns across three diverse lineages. Although this focal measure is symmetric, the underlying novel measure we computed herein, DMS-EX, has several variants that we also considered. Since this measure is based on publicly available deep mutational scanning data, it allows for asymmetric exchangeability estimates, as well as di|ering exchangeabilities between buried and exposed regions, as used in earlier work (Yampolsky and Stoltzfus 2005).

Our motivation was not to produce exchangeability measures that are most useful for every molecular evolution application. Although many of our results focus on comparing which exchangeability measures result in the top-ranked codon models, our goal here was to compare the relative utility of the exchangeability measures themselves. This should not be confused with the goal of producing the most general-purpose codon substitution model, which is why here we did not compare to codon models that incorporate a wider set of parameters (such as M7 and M8 in CODEML) that undoubtedly would fit better than the basic M0 model. Similarly, the intended use of our introduced measures is not to produce improved overall protein alignments or phylogenetic trees, as yielded through the LG matrix for instance (Le and Gascuel 2008). Instead, we hope the measures we introduced will be useful for future approaches such as those that build upon d_R_/d_C_ to infer selection efficacy. The benchmarking we conducted is an important step towards this goal.

Although DEX was most generalizable in our codon substitution models, it is important to emphasize that no single amino acid distance measure will be most informative across all proteins and lineages (Miyata et al. 1979; Braun 2018). For instance, transmembrane proteins have very different amino acid substitution profiles compared to others on average (Mokrab et al. 2010) and such sequences were not analyzed in our study. Similarly, due to their differing evolutionary dynamics partitioning protein structural regions in d_N_/d_S_-like frameworks is known to yield improved insights (Dasmeh et al. 2014; Khan et al. 2015; Meyer and Wilke 2015). This last point was the motivation for our analysis of exchangeability between exposed and buried residues. As expected, we found a clear signal of higher exchangeability at exposed sites. This highlights the value of incorporating this information into selection models (Yang and Swanson 2002), ideally while acknowledging variation in conservation within site classes (Meyer and Wilke 2013).

Although this fact has long been appreciated, incorporating structural information explicitly into d_N_/d_S_-like frameworks remains a non-standard workflow. Recent work has also highlighted the value of asymmetric amino acid substitution matrices (Schulze and Lindorff-Larsen 2026). Our allele frequency analysis highlighted the well-known importance of dividing sequences into buried vs. exposed sites (i.e., the binary classification), but asymmetric exchangeabilities within buried and exposed site separately, or based on their weighted average, did not result in a better fit relative to symmetric matrices. This result could have many explanations. It is possible that asymmetric substitution biases are more protein and/or species-specific than symmetric averages and thus are less generalizable. Instead, the DMS datasets we used to build our measures could be too noisy to provide high-confidence asymmetric exchangeability measures, while providing more useful information for bidirectionally averaged data. In addition, although our variant polarization approach was quite conservative, it is possible that mis-polarizations could erode the asymmetric signal in the allele frequency data. In any case, although exploring asymmetric measures warrants more investigation in codon substitution models, symmetric exchangeability measures are most convenient since standard models typically require time reversible assumptions.

Sophisticated mutation-selection models have previously been developed that allow site-specific amino acid preferences to be fit (Rodrigue et al. 2010; Bloom 2014). Such methods have previously been applied to estimate varying effective population size across lineages (Latrille et al. 2021). These are valuable methods that warrant further application but they suffer from key limitations: inferring amino acid preferences per site is highly computationally intensive and requires extremely deep alignments. Moreover, even with deep alignments, most alignment columns usually will not include instances of most amino acids, meaning that inferring all preferences will not be possible. We see more simplistic measures that build upon d_R_/d_C_, which can allow for faster inferences that are feasible with typical datasets, as complementary to such models.

We hypothesize that a future framework that builds upon our work could investigate questions analogous to those explored in works that compared d_R_/d_C_ across lineages with differing effective population sizes (Weber et al. 2014; Weber and Whelan 2019). In these analyses, some lineages with higher effective population sizes displayed lower d_R_/d_C_ values, which is consistent with more effective selection against radical non-synonymous substitutions (James and Lascoux 2025). An important caveat of these d_R_/d_C_ approaches is that signals could also be explained by shifting amino acid preferences across lineages. More work is needed to determine how to disentangle shifting amino acid preferences from differential efficacy of selection against deleterious non-synonymous substitutions. The amino acid exchangeability measures we explored herein could serve as a basis for such future explorations. Specifically, we believe that the early d_N_/d_S_ method that incorporated amino acid dissimilarity (Goldman and Yang 1994) has been underexplored and could form the basis of revitalized investigation of dR/dC approaches based on improved continuous amino acid exchangeability measures such as DEX. Advances in codon substitution modelling could also be integrated into this framework for potentially improved selection inferences, such as modelling key parameters as distributions (Yang et al. 2000; Murrell et al. 2013) and including protein structural information (Yang and Swanson 2002; Scherrer et al. 2012).

DEX could also be useful as a static comparison for future variant effect predictors. As we and others have highlighted (James and Lascoux 2025), simply knowing which amino acids are involved in a substitution provides substantial insight into how deleterious this substitution is likely to be. However, site-specific context provides additional information (**Supplementary Figures 5 and 6**), and readers should not mistakenly conclude that static distance measures are equally useful as specialized methods, such as RaSP and VespaG, for predicting amino acid substitution impacts. Nonetheless, when evaluating such tools, it is important to compare them to the best possible inference based on amino acid distances alone. Comparing deep learning approaches for predicting substitution impacts to less informative measures, such as Grantham’s or BLOSUM62 distances, could provide misleading confidence in how well a predictor is performing.

In summary, we have compared numerous amino acid distance measures, which vary on a continuum from those based on empirical observations to those based on physicochemical properties. DEX and DMS-EX are the most generalizable and informative of those we tested. These measures will be useful for future work aiming at inferring the efficacy of selection in molecular evolution studies.

## Acknowledgements

We would like to thank members of the North Carolina State Evolutionary Genetics discussion group, Kasie Raymann, and all members of the Bobay Lab for helpful discussions and feedback. We would also like to thank Burkhard Rost, Céline Marquet, Konstantin Weißenow, and Robert Schmirler for their guidance regarding VespaG and related methods. GMD was supported by a Banting Postdoctoral Fellowship, provided by the Canadian government and LMB is supported by the US National Institutes of Health NIGMS (R01GM132137).

## Conflict of interest disclosure

The authors have no conflicts of interest with the content of this article.

## Code and data availability

Code for running all our analyses is available at this GitHub repository: https://github.com/gavinmdouglas/aa_distance_explore. Key datafiles, including the different amino acid distance and similarity metrics compared herein, are available on Zenodo: https://doi.org/10.5281/zenodo.18927704.

## Notes

https://doi.org/10.5281/zenodo.18927704

https://github.com/gavinmdouglas/aa_distance_explore

### Competing Interest Statement

The authors have declared no competing interest.

### Summary of Updates

Major revisions -- 1) The computation of DMS-EX was corrected. The robust Z-scores were previously computed separately at exposed and buried sites before taking the weighted average; they are now computed as an overall robust Z-score (preserving the information that buried sites have lower exchangeability in general) prior to computing the weighting. Both DMS-EX and DEX perform slightly better as a result. 2) Relatedly, the concluding analyses now focus more on buried/exposed and asymmetric exchangeabilities, which represents a more useful application of the allele frequency analyses. The original comparison to the ML approaches has been removed, in agreement with reviewers' comments. These methods are now raised as a discussion point instead, since framing the analysis around how well they predict average exchangeabilities would be misleading given that this is not their intended purpose. 3) A new E. coli dataset focused on full proteins (rather than fragments, as used previously) was substituted in. This is the Alleleome dataset, which provides readily parsed tables. Because it is based on full proteins, obtaining per-residue buried/exposed information is considerably more straightforward. 4) The allele frequency data were polarized by comparison to outgroup sequences in custom alignments. This reduces the number of unambiguous sites available for analysis but allows the unfolded allele frequency spectrum to be used. 5) The structural ML tool ThermoMPNN was added. 6) The title was changed to reference the other measures (as DMS-EX now performs better and is the focus of the allele frequency analysis) and to emphasize benchmarking 7) The model selection criterion was switched from AIC to BIC. This does not change the results, particularly as the number of parameters is the same in most cases, but avoids potential confusion. 8) JTT, WAG, and LG were added. The results are unchanged, but this addresses a common reviewer comment.

