## Supplementary Materials for "DEX: an amino acid exchangeability measure for codon substitution modelling and selection inference"

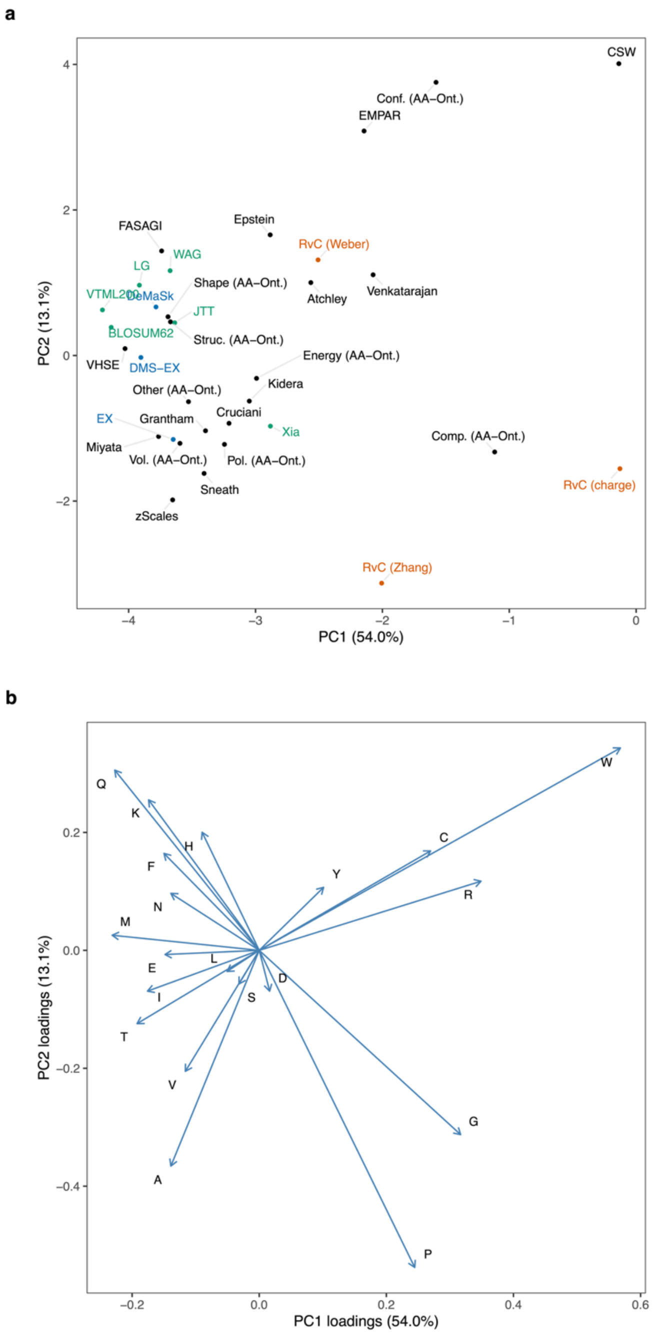


**Supplementary Figure 1**: Principal Components Analysis of all 30 compared measures, based on Euclidean distances between mean exchangeabilities per amino acid. (a) First two principal components (PCs), with measures indicated. See Methods section for all acronyms. (b) Same analysis as in panel a but displaying the loadings that each amino acid contributes to the separation. Percentage of variance explained by each PC is indicated in parentheses. Measure grouping colours match those in Figure 1.

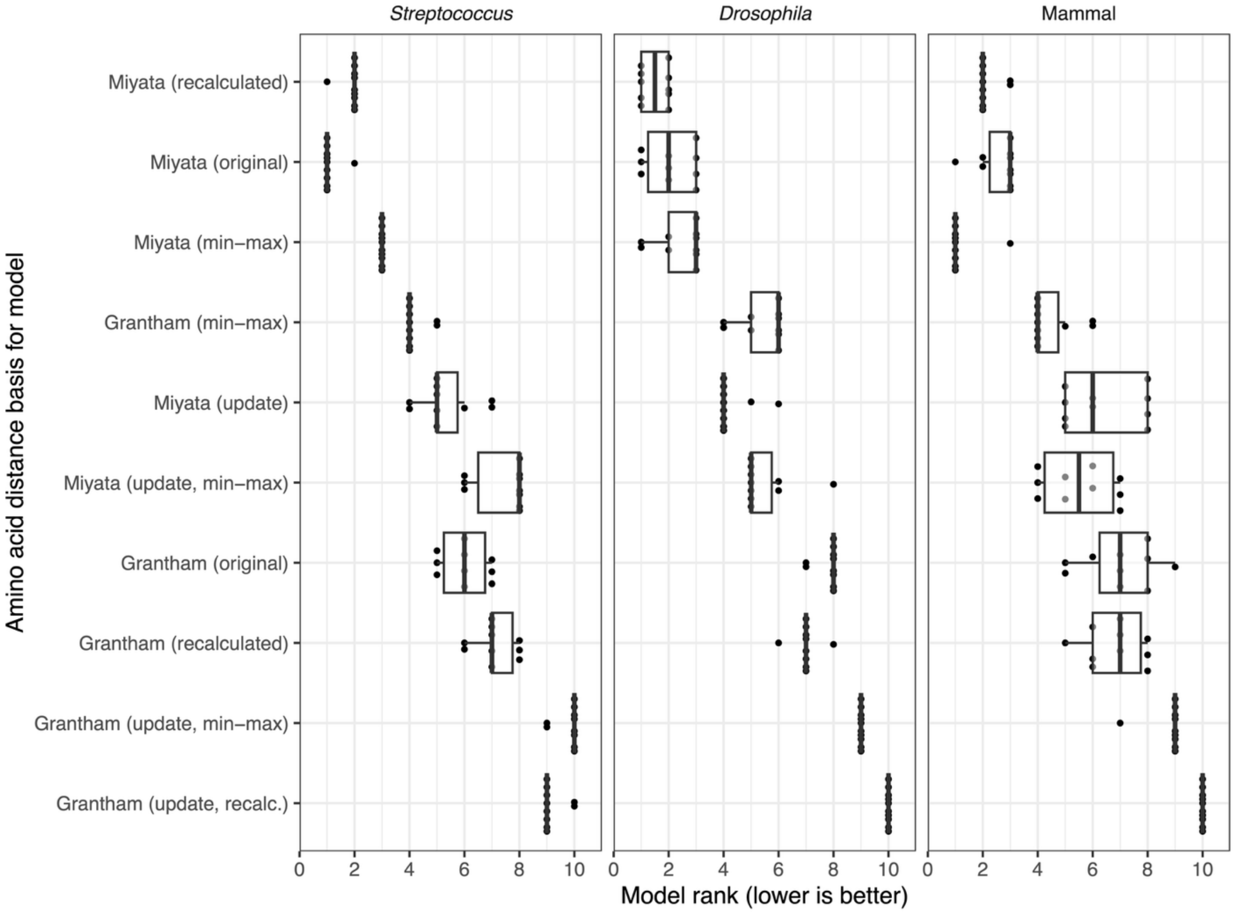


**Supplementary Figure 2**: Model rank results based on Bayesian Information Criterion values, as in main text figures, but for variants of Grantham's and Miyata's amino acid distances.

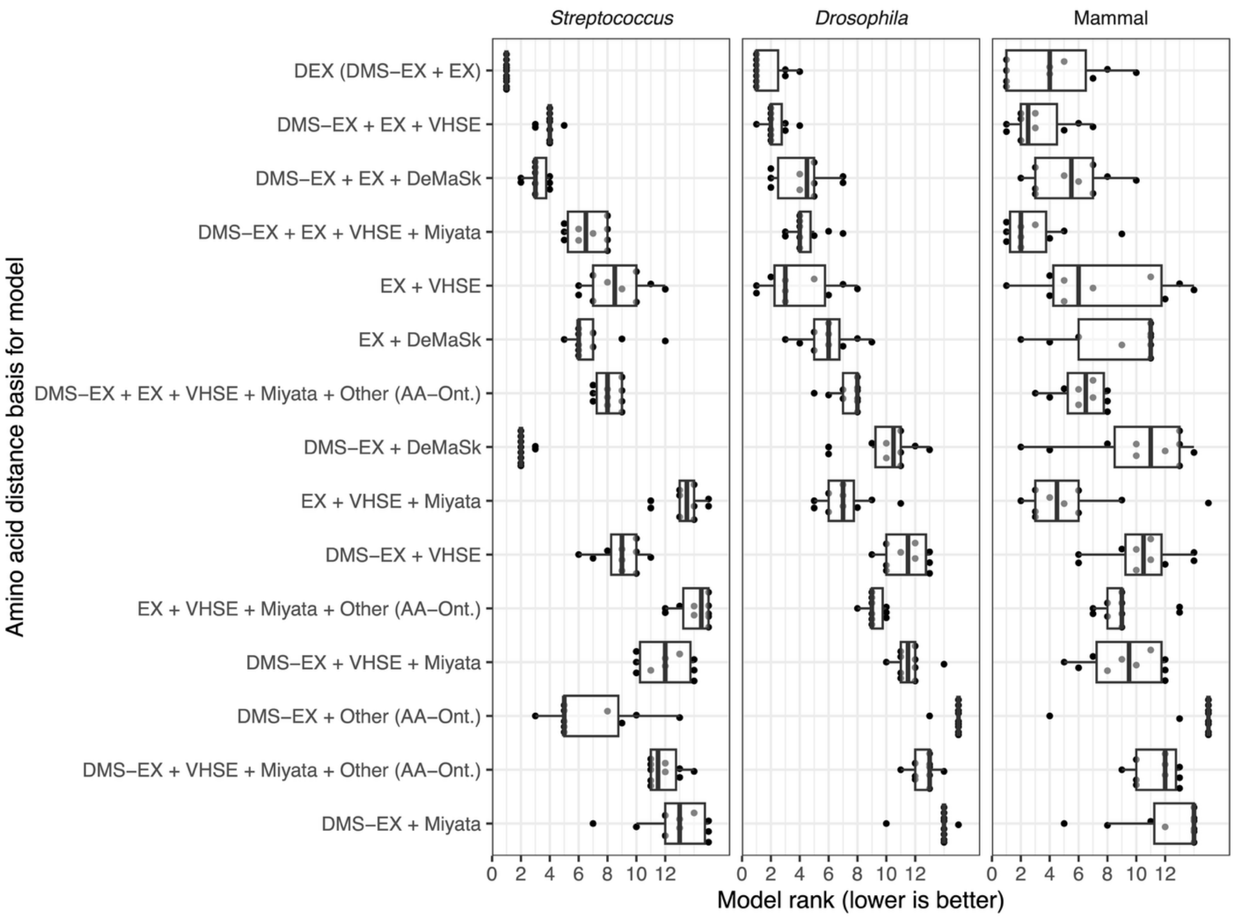


**Supplementary Figure 3**: Model rank results based on Bayesian Information Criterion values but for combined amino acid distance measures produced with the DISTATIS approach. This analysis was conducted on an independent set of 10 alignments compared to Figure 2 (i.e., the same set as displayed in Figure 3).

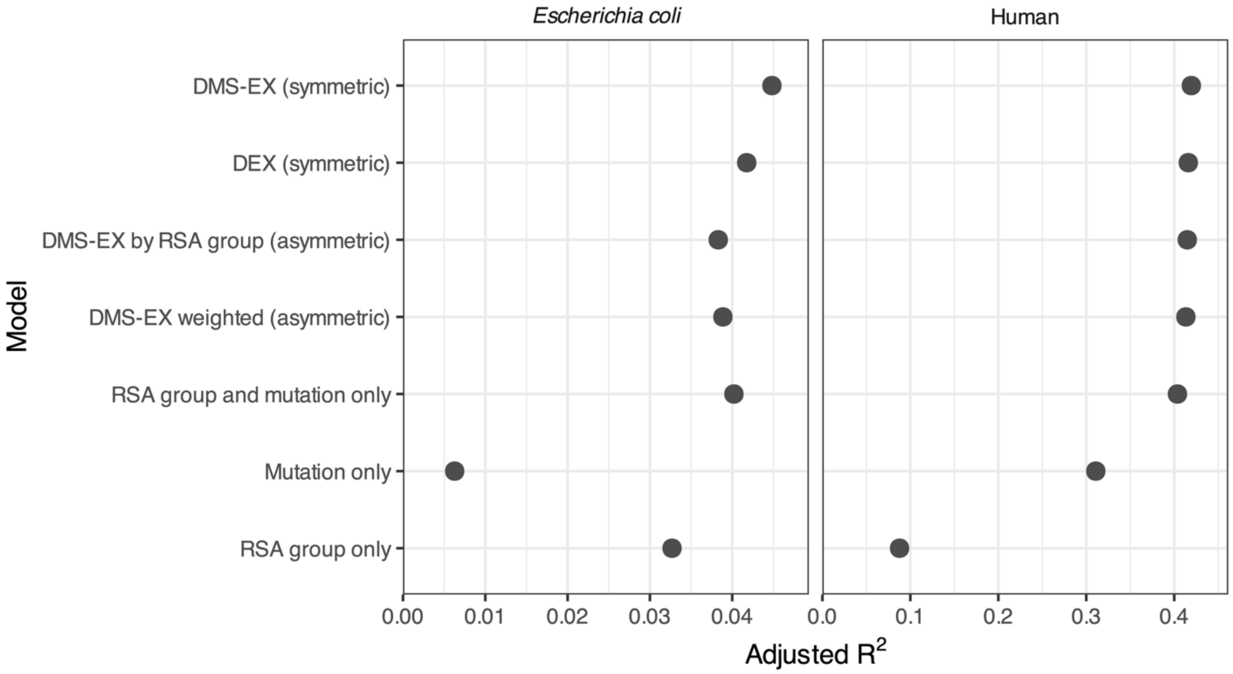


**Supplementary Figure 4**: Adjusted $R^2$ values for linear models predicting proportion of rare segregating amino acid replacements across *E. coli* and human populations. The symmetric DEX and DMS-EX measures match the focal metrics used in the earlier codon substitution models. The models with RSA and/or mutational coefficients only are included to provide baselines over which exchangeability matrices provide an improvement.

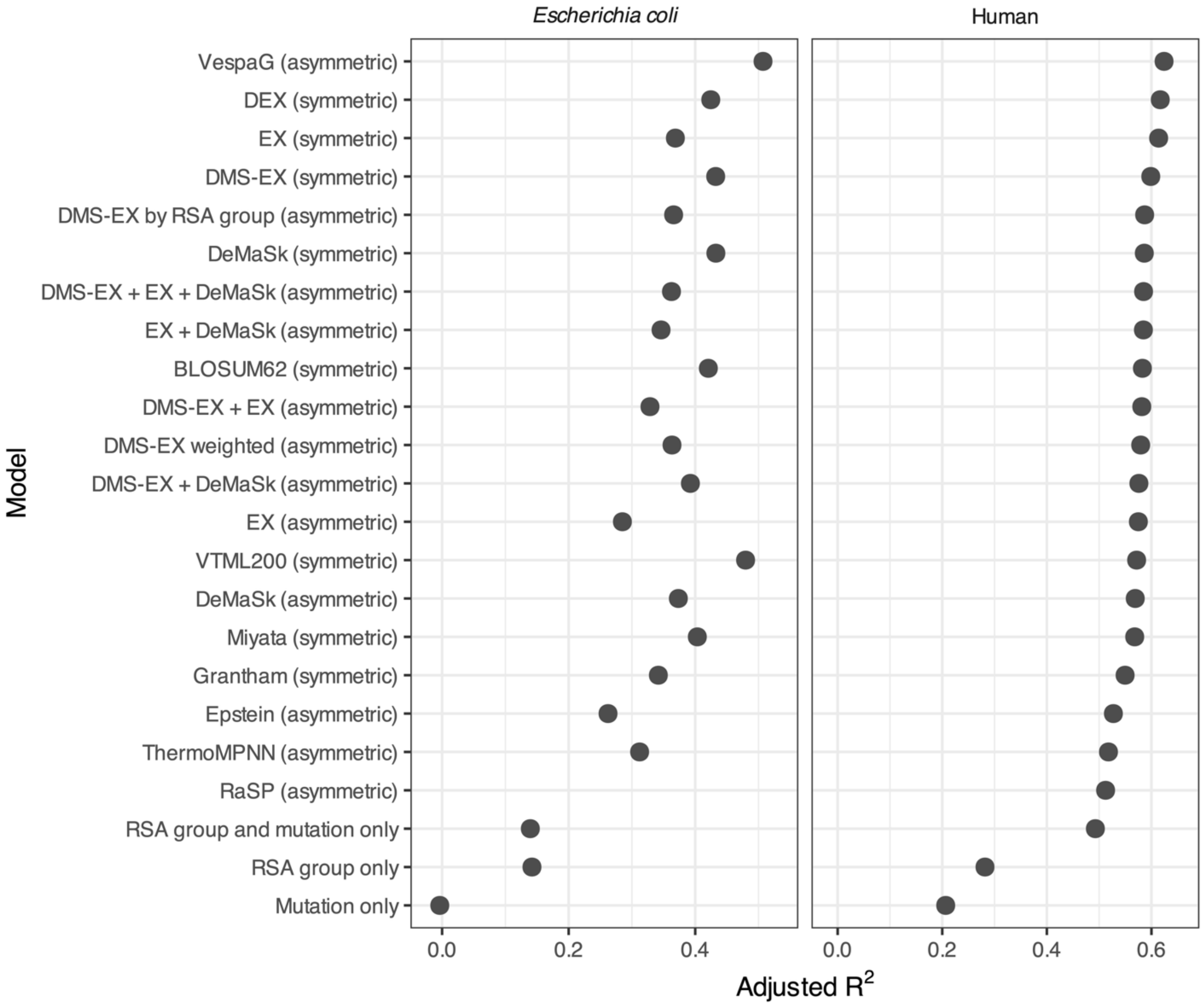


**Supplementary Figure 5**: Adjusted $R^2$ values for linear models predicting mean allele frequencies of amino acid replacements across *E. coli* and human populations, comparing a broader set of measures. The symmetric DEX and DMS-EX measures match the focal metrics used in the earlier codon substitution models. The models with RSA and/or mutational coefficients only are included to provide baselines over which exchangeability matrices provide an improvement.

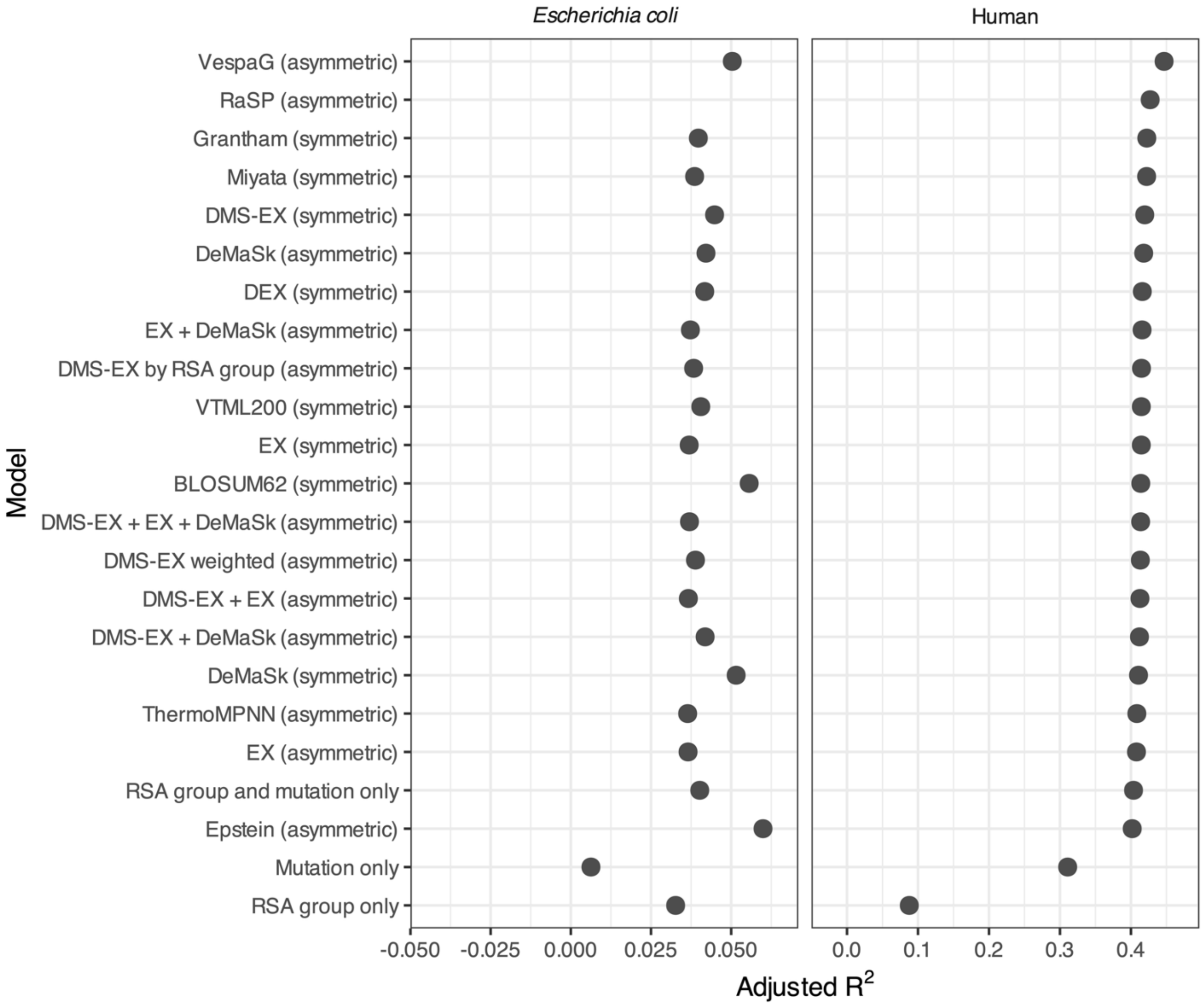


**Supplementary Figure 6**: Adjusted $R^2$ values for linear models predicting the proportion of rare amino acid replacements across *E. coli* and human populations, comparing a broader set of measures. The symmetric DEX and DMS-EX measures match the focal metrics used in the earlier codon substitution models. The models with RSA and/or mutational coefficients only are included to provide baselines over which exchangeability matrices provide an improvement.

**Supplemental Table 1**: Deep mutational scanning (DMS) datasets from the ProteinGym database used to define our DMS-EX measure

| ProteinGym DMS dataset ID | Source organism | Protein name | Selection assay | Article DOI |
|---|---|---|---|---|
| **PTEN_HUMAN_Mighell_2018** | *Homo sapiens* | Phosphatidylinositol 3,4,5-trisphosphate 3-phosphatase and dual-specificity protein phosphatase PTEN | Growth | 10.1016/j.ajhg.2018.03.018 |
| **MSH2_HUMAN_Jia_2020** | *Homo sapiens* | DNA mismatch repair protein Msh2 | Drug resistance | 10.1016/j.ajhg.2020.12.003 |
| **MLAC_ECOLI_MacRae_2023** | *Escherichia coli* | Intermembrane phospholipid transport system binding protein MlaC | Cell growth | 10.1016/j.jbc.2023.104744 |
| **PHOT_CHLRE_Chen_2023** | *Chlamydomonas reinhardtii* | Phototropin | Fluorescence | 10.1021/acssynbio.2c00662 |
| **LGK_LIPST_Klesmith_2015** | *Lipomyces starkeyi* | Levoglucosan kinase | Growth | 10.1021/acssynbio.5b00131 |
| **AMIE_PSEAE_Wrenbeck_2017** | *Pseudomonas aeruginosa* | Aliphatic amidase | Enzyme function | 10.1038/ncomms15695 |
| **PPARG_HUMAN_Majithia_2016** | *Homo sapiens* | Peroxisome proliferator-activated receptor gamma | Expression of CD36 | 10.1038/ng.3700 |
| **CBPA2_HUMAN_Tsuboyama_2023_1O6X** | *Homo sapiens* | Carboxypeptidase A2 | Stability | 10.1038/s41586-023-06328-6 |
| **C6KNH7_9INFA_Lee_2018** | Influenza A virus (A/Perth/16/2009 (H3N2)) | Hemagglutinin | Viral replication | 10.1073/pnas.1806133115 |
| **RNC_ECOLI_Weeks_2023** | *Escherichia coli* | Ribonuclease 3 | Fluorescence | 10.1093/molbev/msad047 |
| **KKA2_KLEPN_Melnikov_2014** | *Klebsiella pneumoniae* | Aminoglycoside 3'-phosphotransferase | Growth | 10.1093/nar/gku511 |
| **OXDA_RHOTO_Vanella_2023_activity** | *Rhodotorula gracilis* | D-amino-acid oxidase | Fluorescence | 10.1101/2023.02.24.529916 |
| **POLG_DEN26_Suphatrakul_2023** | Dengue virus type 2 (strain Thailand/16681/1984) (DENV-2) | Flavivirus NS5 | Viral replication | 10.1101/2023.03.07.531617 |
| **PRKN_HUMAN_Clausen_2023** | *Homo sapiens* | E3 ubiquitin-protein ligase parkin | Protein stability | 10.1101/2023.06.08.544160 |
| **MET_HUMAN_Estevam_2023** | *Homo sapiens* | Hepatocyte growth factor receptor | Cell growth | 10.1101/2023.08.03.551866 |
| **UBC9_HUMAN_Weile_2017** | *Homo sapiens* | SUMO-conjugating enzyme UBC9 | Yeast growth | 10.15252/msb.20177908 |
| **A4D664_9INFA_Soh_2019** | Influenza A virus (A/green-winged teal/Ohio/175/1986 (H2N1)) | Polymerase basic protein 2 | Viral replication | 10.7554/eLife.45079 |
| **A4GRB6_PSEAI_Chen_2020** | *Pseudomonas aeruginosa* | Beta-lactamase VIM-2 | Drug resistance | 10.7554/eLife.56707 |

*This information was lightly modified from the ProteinGym database metadata provided for all datasets. Note that the DMS dataset ID includes the first author's name and year the data was available, which in several cases is the preprint release year rather than the publication year.*

**Supplemental Table 2**: Amino acid dissimilarity/similarity measures compared in this study

| Measure | Category | Brief description | Reference (in main text) |
|---|---|---|---|
| **EX** | Experimental | Exchangeability based on functional impacts of induced substitutions across 12 proteins. | Yampolsky & Stoltzfus 2005 |
| **DMS-EX** | Experimental | New measure of experimental exchangeability computed from deep mutational scanning (DMS) datasets. Not to be confused with the best-performing measure we produced, DEX. | This study |
| **DEX** | Experimental | DISTATIS-based consensus of EX and DMS-EX; top-performing measure overall in our study. | This study |
| **DeMaSk** | Experimental | Exchangeability based on rank-transformed substitution impacts across DMS datasets. | Munro & Singh 2021 |
| **BLOSUM62** | Substitution-based | Log-odds substitution matrix based on observed substitutions between sequences with ≤62% identity. As for all the substitution-based approaches below, it is confounded with mutational biases. | Henikoff & Henikoff 1992 |
| **VTML200** | Substitution-based | Substitution matrix derived from a maximum-likelihood model of evolutionary divergence. | Müller et al. 2002 |
| **JTT** | Substitution-based | Early amino acid substitution rate matrix derived from observed substitutions. | Jones et al. 1992 |
| **WAG** | Substitution-based | Similar to JTT, but based on larger dataset and estimated using maximum likelihood. | Whelan and Goldman 2001 |
| **LG** | Substitution-based | Similar to WAG, but again based on a larger dataset and relaxed certain assumptions. | Le and Gascuel 2008 |
| **Xia** | Substitution-based | Distance based on observed amino acid neighbour frequencies in protein sequences. | Xia & Xie 2002 |
| **RvC (Zhang)** | Radical vs. conservative | Binary classification of substitutions as radical or conservative based on polarity and volume differences. | Zhang 2000 |
| **RvC (Weber)** | Radical vs. conservative | Binary classification of substitutions as radical or conservative based on polarity and volume differences, but with distinct groupings from the above. | Weber & Whelan 2019 |
| **RvC (charge)** | Radical vs. conservative | Binary classification based on charge differences between amino acids. | Zhang 2000 |
| **Grantham** | Physicochemical/structural | Distance based on composition, polarity, and molecular volume. | Grantham 1974 |
| **Miyata** | Physicochemical/structural | Distance based on polarity and molecular volume. | Miyata et al. 1979 |
| **Sneath** | Physicochemical/structural | Proportion of 134 physicochemical characteristics not shared between two amino acids. | Sneath 1966 |
| **Epstein** | Physicochemical/structural | Coefficient of difference based on polarity and size. | Epstein 1967 |
| **EMPAR** | Physicochemical/structural | Dissimilarity based on topological and physicochemical features. | Rao 1987 |

| Measure | Category | Brief description | Reference (in main text) |
|---|---|---|---|
| **CSW** | Physicochemical/structural | Similarity based on pairwise comparisons of backbone dihedral angle distributions across 102 crystal structures. | Kolaskar & Kulkarni-Kale 1992 |
| **Cruciani** | Physicochemical/structural | Distance derived from three scaled principal component scores. | Cruciani et al. 2004 |
| **FASAGI** | Physicochemical/structural | Distance based on six components representing diverse amino acid characteristics. | Liang & Li 2008 |
| **Kidera** | Physicochemical/structural | Distance derived from ten orthogonal factors based on multivariate analysis of physicochemical properties. | Kidera et al. 1985 |
| **zScales** | Physicochemical/structural | Distance from five factors based on physicochemical observations including chromatography and nuclear magnetic resonance data. | Sandberg et al. 1998 |
| **VHSE** | Physicochemical/structural | Distance from eight principal components of hydrophobic, steric, and electronic properties. | Mei et al. 2005 |
| **Atchley** | Physicochemical/structural | Distance from five scaled factors summarizing 54 amino acid features. | Atchley et al. 2005 |
| **Venkatarajan** | Physicochemical/structural | Distance derived from five factors representing 237 physicochemical properties. | Venkatarajan & Braun 2001 |
| **Vol. (AA-Ont.)** | Physicochemical/structural | Distance derived from principal coordinates of volume-related scales in AAontology. | Breimann et al. 2024 |
| **Pol. (AA-Ont.)** | Physicochemical/structural | Distance derived from principal coordinates of polarity-related scales in AAontology. | Breimann et al. 2024 |
| **Struc. (AA-Ont.)** | Physicochemical/structural | Distance derived from principal coordinates of structure/activity-related scales in AAontology. | Breimann et al. 2024 |
| **Energy (AA-Ont.)** | Physicochemical/structural | Distance derived from principal coordinates of energy-related scales in AAontology. | Breimann et al. 2024 |
| **Conf. (AA-Ont.)** | Physicochemical/structural | Distance derived from principal coordinates of conformation-related scales in AAontology. | Breimann et al. 2024 |
| **Shape (AA-Ont.)** | Physicochemical/structural | Distance derived from principal coordinates of shape-related scales in AAontology. | Breimann et al. 2024 |
| **Comp. (AA-Ont.)** | Physicochemical/structural | Distance derived from principal coordinates of composition-related scales in AAontology. | Breimann et al. 2024 |
| **Other (AA-Ont.)** | Physicochemical/structural | Distance derived from principal coordinates of miscellaneous scales in AAontology. | Breimann et al. 2024 |
